# Background selection and the statistics of population differentiation: consequences for detecting local adaptation

**DOI:** 10.1101/326256

**Authors:** Remi Matthey-Doret, Michael C. Whitlock

## Abstract

Background selection is a process whereby recurrent deleterious mutations cause a decrease in the effective population size and genetic diversity at linked loci. Several authors have suggested that variation in the intensity of background selection could cause variation in *F*_*ST*_ across the genome, which could confound signals of local adaptation in genome scans. We performed realistic simulations of DNA sequences, using parameter estimates from humans and sticklebacks, to investigate how variation in the intensity of background selection affects different statistics of population differentiation. We show that, in populations connected by gene flow, Weir & Cockerham’s (1984) estimator of *F*_*ST*_ is largely insensitive to locus-to-locus variation in the intensity of background selection. Unlike *F*_*ST*_, however, *d*_*XY*_ is negatively correlated with background selection. We also show that background selection does not greatly affect the false positive rate in *F*_*ST*_ outlier studies. Overall, our study indicates that background selection will not greatly interfere with finding the variants responsible for local adaptation.

## Introduction

Natural selection affects patterns of genetic diversity throughout the genome. How selection affects genetic diversity on a single isolated locus is relatively easy to model; however, when a large number of linked loci are considered, interactions between evolutionary pressures at different sites render the task of modelling much more difficult.

Maynard Smith & Haigh (1974) recognized the influence of selection on linked neutral sites, proposing that strong positive selection could reduce genetic diversity at nearby sites. This process is now referred to as a ‘selective sweep’. Much later, Charlesworth et al. (1993) proposed that deleterious mutations could also affect genetic diversity at nearby sites, because some haplotypes would be removed from the population as selection acts against linked deleterious alleles. They named this process background selection (BGS). Both selective sweeps and background selection affect genetic diversity; they both reduce the effective population size of linked loci. Empirical evidence of a positive correlation between genetic diversity and recombination rate has been reported in several species (Cutter and Payseur, 2013), including *Drosophila melanogaster* (Begun & Aquadro, 1992; Elyashiv et al., 2016), humans (Spencer et al., 2006), collared flycatchers, hooded crows and Darwin’s finches (Dutoit et al., 2017; see also Vijay et al., 2017).

BGS is also expected to affect *F*_*ST*_ (Charlesworth et al., 1997; Cutter & Payseur, 2013; Cruickshank & Hahn, 2014; Hoban et al., 2016). At low effective population size, different populations may randomly fix different alleles, increasing *F*_*ST*_, while at high population size, allele frequency changes less through time and different populations are more likely to have comparable allele frequencies, keeping *F*_*ST*_ low. This negative relationship between effective population size *N*_*e*_ and *F*_*ST*_ is captured in Wright’s classical infinite island result; 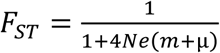 (Wright, 1943). One might therefore expect that loci under stronger BGS would show higher *F*_*ST*_.

Many authors have also argued that, because BGS reduces the within-population diversity, it should lead to high *F*_*ST*_ (Cutter & Payseur, 2013; Cruickshank & Hahn, 2014; Hoban et al., 2016). Expressed in terms of heterozygosities, 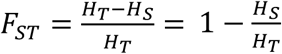, where *H*_*T*_ is the expected heterozygosity in the entire population and *H*_*S*_ is the average expected heterozygosity within subpopulations (*H*_*S*_ and *H*_*T*_ are also sometimes called π_S_ and π_T_; e.g. Charlesworth, 1998). All else being equal, a decrease of *H*_*S*_ would indeed lead to an increase of *F*_*ST*_. However, all else is not equal; *H*_*T*_ is also affected by BGS (Charlesworth et al., 1997). Therefore in order to understand the effects of BGS on *F*_*ST*_, we must understand the relative impact of BGS on both *H*_*S*_ and *H*_*T*_.

Performing numerical simulations, Charlesworth et al. (1997) report that BGS reduces the within population heterozygosity *H*_*S*_ slightly more than it reduces the total heterozygosity *H*_*T*_, causing a net increase in *F*_*ST*_. The effect on *F*_*ST*_ reported is quite substantial, but, importantly, their simulations were not meant to be realistic. The authors highlighted their goal in the methods:

> “The simulations were intended to show the qualitative effects of the various forces studied […], so we did not choose biologically plausible values […]. Rather, we used values that would produce clear-cut effects”.

For example, talking about their choice for the deleterious mutation rate of 8 × 10^−4^ per site:

> “This unrealistically high value was used in order for background selection to produce large effects […]”

Much of the literature on the effect of BGS on *F*_*ST*_ is based on the results in Charlesworth et al. (1997), even though they only intended to show proof of concept (see also Zeng & Charlesworth, 2011 and Zeng & Corcoran, 2015). They did not attempt to estimate how strong of an effect BGS has on *F*_*ST*_ in real genomes.

It is important to distinguish two separate questions when discussing the effect of BGS on *F*_*ST*_; 1) How does BGS affect the average genome-wide *F*_*ST*_? and 2) How does locus-to-locus variation in the intensity of BGS affect locus-to-locus variation in *F*_*ST*_? The second question is of particular interest to those trying to identify loci under positive selection (local selection or selective sweep). Locus-to-locus variation in *F*_*ST*_ potentially could be confounded with the *F*_*ST*_ peaks created by positive selection. In this paper, we focus on this second question.

The identification of loci involved in local adaptation is often performed via *F*_*ST*_ outlier tests (Lotterhos & Whitlock, 2014; Hoban et al., 2016). Other tests exist to identify highly divergent loci such as cross-population extended haplotype homozygosity (XP-EHH; Sabeti et al., 2007), comparative haplotype identity (Lange & Pool, 2016), cross-population composite likelihood ratio (XP-CLR; Chen et al., 2010). *F*_*ST*_ outlier tests, such as FDist2 (Beaumont & Nichols, 1996), BayeScan (Foll & Gaggiotti, 2008) or FLK (Bonhomme et al., 2010), look for genomic regions showing particularly high *F*_*ST*_ values to find candidates for local adaptation. If BGS can affect *F*_*ST*_ unevenly across the genome, then regions with a high intensity of BGS could potentially have high *F*_*ST*_ values that could be confounded with the pattern caused by local selection (Charlesworth et al., 1997; Cruickshank & Hahn, 2014). BGS could therefore inflate the false positive rate when trying to detect loci under local selection.

The potential confounding effect of BGS on signals of local adaptation has led to an intense effort trying to find solutions to this problem (Bank et al., 2014; Huber et al., 2016). Many authors have understood from Cruickshank and Hahn (2014) that *d*_*XY*_ should be used instead *F*_*ST*_ in outlier tests (e.g. McGee et al., 2015; Yeaman, 2015; Whitlock & Lotterhos, 2015; Brousseau et al., 2016; Picq et al., 2016; Payseur & Rieseberg, 2016; Hoban et al., 2016; Vijay et al., 2017; see also Nachman & Payseur, 2012). *F*_*ST*_ is a measure of population divergence relative to the total genetic diversity, while *d*_*XY*_ is an absolute measure of population divergence defined as the probability of non-identity by descent of two alleles drawn in the two different populations averaged over all loci (Nei, 1987; Nei, 1987 originally called it *D*_*XY*_ but, here, we follow Cruickshank and Hahn’s, 2014 terminology by calling it *d*_*XY*_). The argument is that because *F*_*ST*_ is a measure of divergence relative to the genetic diversity and *d*_*XY*_ an absolute measure of divergence and because BGS reduces genetic diversity, then BGS must affect *F*_*ST*_ but not *d*_*XY*_, a claim that we will investigate in this paper.

Whether BGS can affect genome-wide *F*_*ST*_ under some conditions is not in doubt (Charlesworth et al., 1997), but whether locus-to-locus variation in the intensity of BGS present in natural populations substantially affects variation in *F*_*ST*_ throughout the genome is very much unknown. Empirically speaking, it has been very difficult to measure how much of the genome-wide variation in genetic diversity is caused by BGS, as opposed to selective sweeps or variation in mutation rates (Cutter & Payseur, 2013; see also attempts in humans by Cai et al., 2009 and McVicker et al. 2009). We are therefore in need of realistic simulations that can give us more insight into how BGS affects genetic diversity among populations and how it affects the statistics of population divergence.

In this article, we investigate the effect of BGS in structured populations with realistic numerical simulations using parameter estimates from humans and stickleback. Our two main goals are 1) to quantify the impact of locus-to-locus variation in the intensity of BGS on *d*_*XY*_ (Nei, 1987) and *F*_*ST*_ (Weir & Cockerham, 1984) and 2) to determine whether BGS inflates the false positive rate of *F*_*ST*_ outlier tests.

## Methods

Our goal is to perform biologically plausible simulations of the local genomic effects of background selection. BGS is expected to vary with gene density, mutation rate and recombination rate across the genome. We used data from real genomes to simulate realistic covariation in recombination rates and gene densities. We chose to base our simulated genomes on two eukaryote genomes, sticklebacks and humans, because these two species have attracted a lot attention in studies of local adaptation and because sticklebacks have a variance in recombination rate which is almost 15 times higher than humans (data not shown), allowing us to test vastly different types of eukaryotic genomes. The recombination rate variation in humans is extremely fine scale, but it presents the potential issue that it is estimated from linkage disequilibrium data. As selection causes linkage disequilibrium to increase, estimates of recombination rate at regions under strong selection may be under-estimated, hence artificially increasing the simulated variance in the intensity of BGS. Although the recombination map for stickleback is much less fine scaled, the estimates are less likely to be biased as they are computed from pedigrees.

Our simulations are forward in time and were performed using the simulation platform SimBit version 3.69. The code and user manual are available at https://github.com/RemiMattheyDoret/SimBit. To double check our results, we also ran some simulations with SFS_code (Hernandez, 2008), confirming that we get consistent distributions of genetic diversity and of *F*_*ST*_ among simulations (results not shown). Generations are non-overlapping, individuals are hermaphrodites, mating is random within patches and selection occurs before dispersal.

### Genetics

For each simulation, we randomly sampled a sequence of about 10 cM coming either from the stickleback (*Gasterosteus aculeatus*) genome or from the human genome (see treatments below) and used this genomic location to determine the recombination map and exon locations for a simulation replicate. For the stickleback genome, we used the gene map and recombination map from Roesti et al. (2013). Ensembl-retrieved gene annotations were obtained from Marius Roesti. For the human genome, we used the recombination map from The International HapMap Consortium (2007) and the gene positions from NCBI and positions of regulatory sequences on Ensembl (Zerbino et al., 2017). We excluded sex chromosomes to avoid complications with haploid parts of the genomes. As estimates of mutation rate variation throughout the genome are very limited, we assumed that the haploid mutation rate varies from site to site following an exponential distribution with mean of 2.5×10^−8^ per generation (Nachman & Crowell, 2000).

More specifically, we first randomly sampled a sequence of 10^5^ nucleotides, which we will refer to as the focal region. All of the statistics (defined under the section *Statistics* below) are calculated only on the focal region of each simulation. Nucleotides that occur in locations determined to be exons in the sampled genomic map are subject to selection (see *Selection*), while all other nucleotides are assumed to be neutral. The focal region itself contained on average ∼0.44 genes for the human genome and ∼3.15 genes for the stickleback genome.

We simulated a 5 cM region on each side of the focal region (resulting in a window of 10 cM plus the recombination rate present in the specific focal region of 10^5^ sites) in order to capture the local effects of background selection. In these 10 cM flanking regions, we only tracked exons. In the nearest 1 cM on each side of the focal region, as well within the focal region, we individually simulated each nucleotide with a biallelic locus. On the remaining outer 4 cM, to improve the speed and RAM usage of the process, we tracked the number of mutations in blocks of up to 100 nucleotides. For these blocks, we tracked only the number of mutations but not their location within the block. Ignoring recombination within a block likely had little effect on the results because the average recombination distance between the first and last site of a block is of the order of 10^−6^ cM. The expected number of segregating sites within a block is 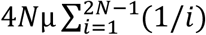, which for a mutation rate per block of 10^−6^ and a population size of *N* = 10,000 is ∼0.42. The probabilities of having more than one mutation and more than two mutations (based on a Poisson approximation) are therefore only approximately 6.7% and 0.9%, respectively. Overall, the level of approximation used is very reasonable.

### Selection

As we are interested in the effect of BGS, we modelled the effects of purifying selection against novel deleterious mutations. Each nucleotide in the exons (and regulatory sequences for the human genome) is subject to purifying selection with a selection coefficient against mutant alleles determined by a gamma distribution described below. For focal regions that include exons, statistics are computed over a sequence that is at least partially under direct purifying selection.

To create variance in selection pressures throughout the genome, each exon (and regulatory sequence for the human genome) has its own gamma distribution of heterozygous selection coefficients *s*. The mean and variance of these gamma distributions are drawn from a bivariate uniform distribution with correlation coefficient of 0.5 (so that when the mean is high, so is the variance) bounded between 10^−8^ and 0.2 for both the mean and the variance. These bounds were inspired by the methodology used in Gilbert et al. (2017). The gamma distributions are bounded to one. Figure S1 shows the overall distribution of selection coefficient *s*, with 2% of mutations being lethal and an average deleterious selection coefficient for the non-lethal mutations of 0.07. To improve the performance of our simulations, we used multiplicative dominance, where the fitness of heterozygotes is at locus *i* is 1-*s*_*i*_ and the fitness of the double mutant is (1-*s*_*i*_) ^2^.

As a consequence of our parameter choices, our genome-wide deleterious mutation rate was about 1.6 in sticklebacks and about 3 in humans. 9.4% of the stickleback genome and 2.6% of the human genome was under purifying selection. For comparison, the genome-wide deleterious mutation rate is estimated at 2.2 in humans (Keightley, 2012) and 0.44 in rodents (Keightley & Gaffney, 2003). To our knowledge, there is currently no such estimation for sticklebacks. Note however that the above estimates cannot reliably detect mutations that are quasi lethal (*s* << 1/2*N*). By our distribution of selective coefficients, 49% of all deleterious mutations have a heterozygote selective coefficient lower than 1/2*N*_*e*_ when *N*_*e*_ = 1,000 (42% when *N*_*e*_ = 10,000).

It is worth noting however that, in rodents, about half of the deleterious mutation rate occurs in non-coding sequences (Keightley & Gaffney, 2003). Our simulations using human genome had all exons and all regulatory sequences under purifying selection. With our simulations based on the stickleback genome, however, only exons were under purifying selection. It is therefore possible that we would have over-estimated the deleterious mutation rate in gene-rich regions and under-estimated the deleterious mutation rate in other regions, especially in stickleback. This would artificially increase the locus-to-locus variation in the intensity of BGS in our simulations.

### Demography

In all simulations, we started with a burn-in phase with a single population of *N* diploid individuals, lasting 5 × 2*N* generations. The population was then split into two populations of *N* individuals each with a migration rate between them equal to *m*. After the burn-in phase, each simulation was run for 5 × 2*N* more generations for a total of 10 × 2*N* generations.

### Treatments

We explored presence and absence of deleterious mutations over two patch sizes, three migration rates, and two genomes.

We do not have a full factorial design. We considered a basic design and explored variations from this design. The basic design had a population size per patch of *N* = 1000, a migration rate of m = 0.005 and used the stickleback genome for its recombination map and gene positions. As deviations from this basic design, we explored modification of every variable, one variable at a time. The *Large N* treatment has N = 10000. The *Human* treatment uses the human genome for gene positions, regulatory sequences and recombination map. The treatments *No Migration* and *High Migration* have migration rates of *m* = 0 and *m* = 0.05, respectively.

To test the robustness of our results and because it may be relevant for inversions, we also had unrealistic simulations where recombination rate for the entire genome was set at zero. As a check against previous work, we qualitatively replicated the results Charlesworth et al. (1997) by performing simulations with similar assumptions as they used. We named this treatment *CNC97*. In our *CNC97* simulations, *N*=2000, *m*=0.001, and 1000 loci were all equally spaced at 0.1 cM apart from each other with constant selection pressure with heterozygotes having fitness of 0.98 and double homozygotes fitness of 0.9 and constant mutation rate μ = 0.0004.

In all treatments (except *Large N*), we performed 4000 simulations; 2000 simulations with BGS and 2000 simulations without selection (where all mutations were neutral). For *Large N*, simulations took more memory and more CPU time. We therefore could only perform 2000 simulations for *Large N*; 1000 simulations with background selection and 1000 simulations without selection. That represents a total of 26,000 simulations for 7 treatments. A full list of all treatments can be found in table 1.

**Table 1:**
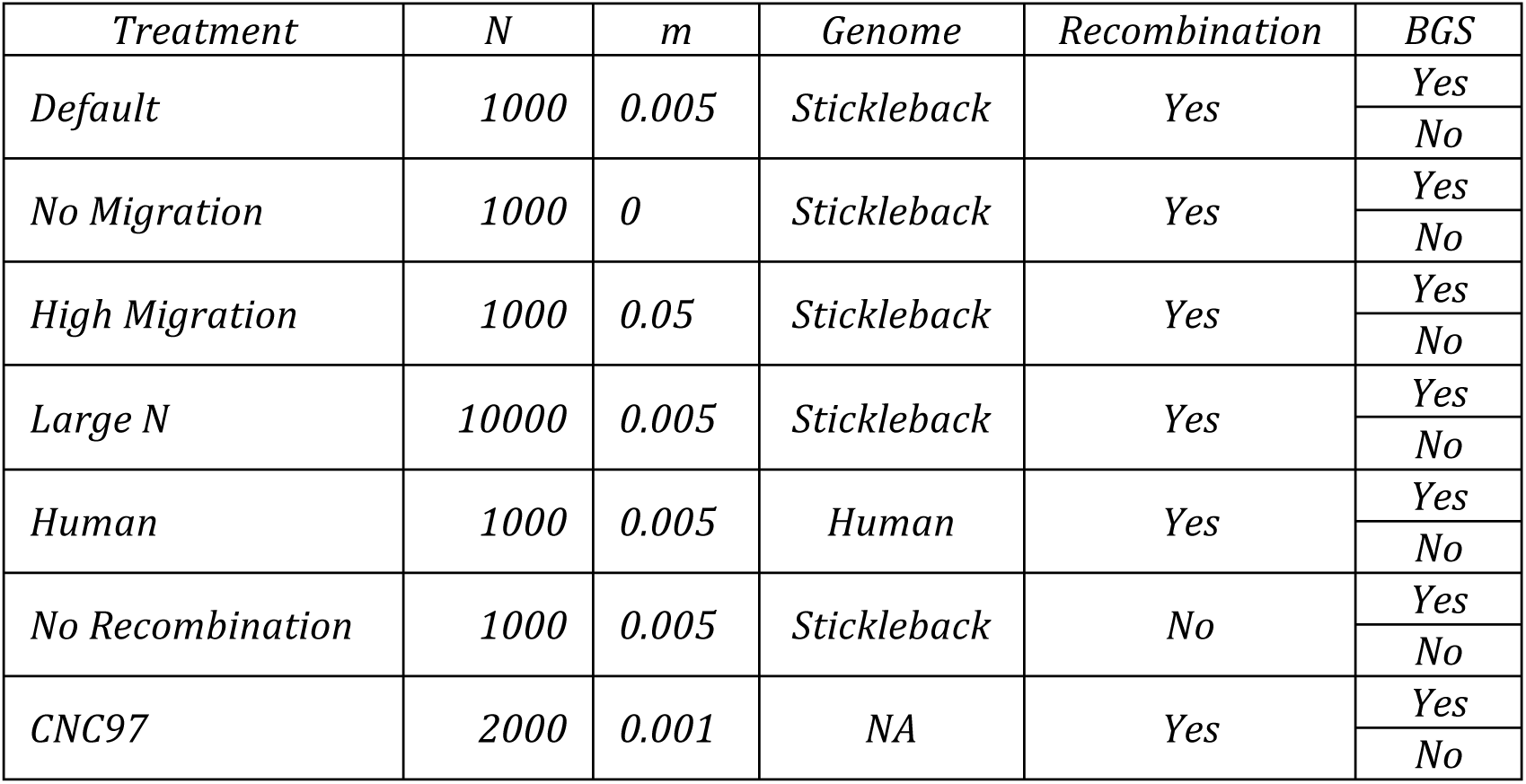
Summary of treatments. For all treatments but *CNC97*, the average mutation rate was set to 2.5 × 10^−8^ per site, per generation and the mean heterozygous selection coefficient to 0.1.

| *Treatment* | *N* | *m* | *Genome* | *Recombination* | *BGS* |
| --- | --- | --- | --- | --- | --- |
| *Default* | *1000* | *0.005* | *Stickleback* | *Yes* | *Yes* |
|  |  |  |  |  | *No* |
| *No Migration* | *1000* | *0* | *Stickleback* | *Yes* | *Yes* |
|  |  |  |  |  | *No* |
| *High Migration* | *1000* | *0.05* | *Stickleback* | *Yes* | *Yes* |
|  |  |  |  |  | *No* |
| *Large N* | *10000* | *0.005* | *Stickleback* | *Yes* | *Yes* |
|  |  |  |  |  | *No* |
| *Human* | *1000* | *0.005* | *Human* | *Yes* | *Yes* |
|  |  |  |  |  | *No* |
| *No Recombination* | *1000* | *0.005* | *Stickleback* | *No* | *Yes* |
|  |  |  |  |  | *No* |
| *CNC97* | *2000* | *0.001* | *NA* | *Yes* | *Yes* |
|  |  |  |  |  | *No* |

We set the generation 0 at the time of the split. The state of each population was recorded at the end of the burn-in period (generation −1) and at generations 0.001× 2*N*, 0.05 × 2*N*, 0.158 × 2*N*, 1.581× 2*N* and 5 × 2*N* after the split. For *N*=1000, the sampled generations are therefore −1, 2, 100, 316, 3162 and 10000.

### Predicted intensity of Background Selection

In order to investigate the locus-to-locus correlation between the predicted intensity of BGS and various statistics, we computed *B*, a statistic that approximates the expected ratio of the coalescent time with background selection over the coalescent time without background selection 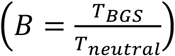. *B* quantifies how strong BGS is expected to be for a given simulation (Nordborg et al. 1996). A *B* value of 0.8 means that BGS has caused a drop of genetic diversity of 20% compared to a theoretical absence of BGS. Lower *B* values indicate stronger BGS.

Both Hudson & Kaplan (1995) and Nordborg et al. (1996) have derived theoretical expectations for *B*. We applied both methods and found that the predictions of the two formulas are highly correlated (Figure S2). Because the Hudson & Kaplan (1995) approach has been more popular in the literature (cited almost twice as often), we show only the *B* values computed following Hudson & Kaplan (1995):

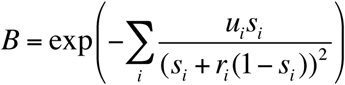

where *r*_*i*_ is the recombination rate between the focal site and the *i*^th^ site under selection, and *s*_*i*_ is the heterozygous selection coefficient at that site. *u*_*i*_ is the mutation rate at the *i*^th^ site. By this formula, *B* is bounded between 0 and 1, where 1 means no BGS at all and low values of *B* mean strong BGS. We computed *B* for all sites in the focal region and report the average *B* for the region.

For the stickleback genome, *B* values ranged from 2 × 10^−6^ to 1.0 with a mean of 0.937 (Figure S2). For the human genome, *B* values ranged from 0.45 to 1.0 with a mean at 0.975. There is indeed less variability and much fewer extremely low *B* values in the human genome. In the unrealistic *No Recombination* treatment, *B* values range from 0.00003 to 0.84 with a mean of 0.17.

### F_ST_ outlier tests

In order to know the effect of BGS on outlier tests of local adaptation, we used a variant of FDist2 (Beaumont & Nichols, 1996). We chose FDist2 because it is a simple and fast method for which the assumptions of the test match well to the demographic scenario simulated here. Because the program FDist2 is not available through the command line, we rewrote the FDist2 algorithm in R and C++. Source code can be found at https://github.com/RemiMattheyDoret/Fdist2.

Our FDist2 procedure is as follows; first, we estimated the migration rate from the average 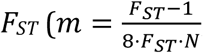; Charlesworth, 1998) and then running 50000 simulations each lasting for 50 times the half-life to reach equilibrium *F*_*ST*_ given the estimated migration rate (Whitlock, 1992). For each SNP, we then selected the subset of FDist2 simulations for which allelic diversity was less than 0.02 away from the allelic diversity of the SNP of interest. The *P*-value is computed as the fraction of FDist2 simulations within this subset having a higher *F*_*ST*_ than the one we observed. The false positive rate is then defined as the fraction of neutral SNPs for which the *P*- value is lower than a given α value. The α values explored are 0.1, 0.05, 0.01, 0.001, 0.0001 and 0.00001.

For the outlier tests, to avoid issues of pseudo-replication, we considered only a single SNP per simulation whose minor allele frequency is greater than 0.05. Then, we randomly assembled SNPs from a given treatment into groups of 500 SNPs to create the data file for FDist2. We have 4000 simulations (2000 with BGS and 2000 without BGS) per treatment (*Large N* is an exception with only 2000 simulations total), which allowed 8 independent false positive rate estimates per treatment (4 estimates with BGS and 4 without BGS). In each treatment, we tested for different false positive rate with and without BGS with both a Welch’s *t*-test and a Wilcoxon test.

### Statistics

F_*ST*_ and *d*_*XY*_ are both measures of population divergence. In the literature there are several definitions of *F*_*ST*_, and we also found potential misunderstanding about how *d*_*XY*_ is computed. We want to clarify here these definitions and what we mean when we use the terms *F*_*ST*_ and *d*_*XY*_.

There are two main estimators of *F*_*ST*_ in the literature; *G*_*ST*_ (Nei, 1973) and θ (Weir & Cockerham, 1984). In this article, we focus on θ as an estimate of *F*_*ST*_ (Weir & Cockerham, 1984). There are also two methods of averaging *F*_*ST*_ over several loci. The first method is to simply take an arithmetic mean over all loci. The second method consists at calculating the sum of the numerator of θ over all loci and dividing it by the sum of the denominator of θ over all loci. Weir and Cockerham (1984) showed that this second averaging approach has lower bias than the simple arithmetic mean. We will refer to the first method as the “average of ratios” and to the second method as “ratio of the averages” (Reynolds et al. 1983; Weir & Cockerham, 1984). In this article, we use *F*_*ST*_ as calculated by “ratio of the averages”, as advised by Weir and Cockerham (1984). To illustrate the effects of BGS on the biased estimator of *F*_*ST*_, we also computed *F*_*ST*_ as a simple arithmetic mean (“average of the ratio”), and we will designate this statistic with a subscript *F*_*ST* (*average of ratios*)_. d_*XY*_ is a measure of genetic divergence between two populations X and Y. Nei (1987) defined *d*_*XY*_ as

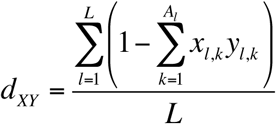

where *L* is the total number of sites, *A*_*l*_ is the number of alleles at the *l*^th^ site and *x*_*l,k*_ and *y*_*l,k*_ are the frequency of the *k*^th^ allele at the *l*^th^ locus in the population X and Y respectively.

Some population genetics software packages (e.g., EggLib; De Mita and Siol, 2012) average *d*_*XY*_ over polymorphic sites only, instead of averaging over all sites, as in Nei’s (1987) original definition of *d*_*XY*_. This measure averaged over polymorphic sites only will be called *d*_*XY-SNP*_; otherwise, we use the original definition of *d*_*XY*_ by Nei (1987).

The statistics reported are the average *F*_*ST*_, *d*_*XY*_, and within population genetic diversity 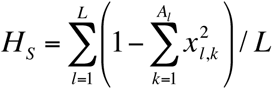. For each treatment and at each generation, we computed five independent Pearson correlation tests between *B* and *F*_*ST*_, *F*_*ST* (*average of* ratios)_, *d*_*XY*_, *d*_*XY-SNP*_ and *H*_*S*_. We compared our correlation tests with ordinary least squares regressions and robust regressions (using M-estimators; Huber, 1964), and the results were consistent.

## Results

Correlations between the statistics *H*_*S*_, *F*_*ST*_, *F*_*ST* (*average of ratios*)_, *d*_*XY*_, and *d*_*XY-SNP*_ and *B*, are summarized in tables S1, S2, S3, S4 and S5, respectively. Figure 1 shows the means and standard errors for the treatments *Default, High Migration, Large N, Human* and *No Migration*. The same graphs for the treatments *No Recombination* and *CNC97* can be found in Figure S3.

**Figure 1:**
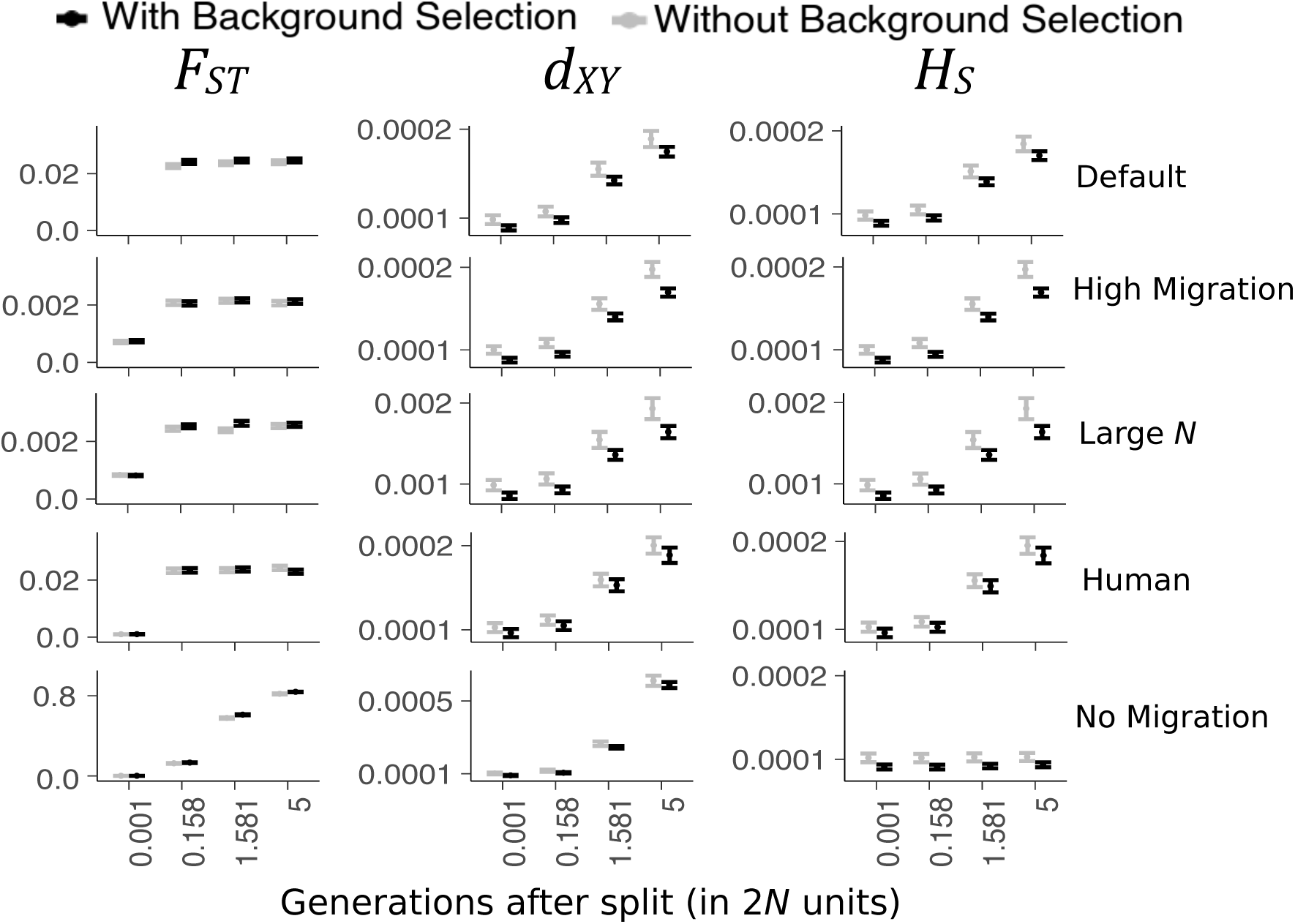
Comparisons of means *F*_*ST*_ (left column), *d*_*XY*_ (central column), and *H*_*S*_ (right column) between simulations with (black) and without (grey) BGS. Similar graphs for treatments *No Recombination* and *CNC97* are in figure S3. Error bars are 95% CI.

### Genetic Diversity

Genetic diversity within populations (*H*_*S*_) is very similar among the treatments *Default, High Migration* and *Human* (around *H*_*S*_ = 1.9 × 10^−4^) but is about 1.9 times lower in the *No Migration* treatment (*H*_*S*_ = 1.0 × 10^−4^) and about 10 times higher in the *Large N* treatment (*H*_*S*_ = 1.9 × 10^−3^; Figure 1).

*B* is significantly correlated with genetic diversity within populations (*H*_*S*_) for all treatments using the stickleback genome (and at almost all generations) but not with the *Human* treatment (table S1). Excluding the unrealistic treatments (*No Recombination* and *CNC97*), simulations with BGS have a genetic diversity 4% to 20% lower than simulations without BGS (Figure 1, right column and Figure S3, right column). In the *Human* treatment, there is no significant correlation of *B* and *H*_*S*_. Note that Pearson’s correlation coefficients between *B* and *H*_*S*_ are always very small even when the effect is highly significant. The largest *R*^2^ observed in realistic simulations (excluding in the *No Recombination* and *CNC97* treatments) between *B* and *H*_*S*_ is *R*^2^ ≈ 0.0121.

### Statistics of population divergence

Figure 2 shows the correlation between *B* and the statistics *F*_*ST*_, *d*_*XY*_ and *H*_*S*_ for *Default* at the last generation. These graphs highlight the general tendencies of the treatments *Default, High Migration, Large N* and *No Recombination*. The strongest correlation with *B* is observed for the statistics *d*_*XY*_ (*P* = 3.28 × 10^−5^, *R* = 0.093) and *H*_*S*_ (*P* = 3.1 × 10^−5^, *R* = 0.093). In fact, the two statistics *d*_*XY*_ and *H*_*S*_ are very highly correlated (*P* < 2.2 × 10^−16^, *R* = 0.99). This high correlation explains the resemblance between the central and right graphs of figure 2. *F*_*ST*_ is not correlated with B (*P* = 0.99, *R* = 10^−4^). All correlation tests between *B* and the statistics *F*_*ST*_, *F*_*ST* (*average of ratios*)_, *d*_*XY*_, *d*_*XY-SNP*_ can be found in Tables S2, S3, S4 and S5, respectively.

**Figure 2:**
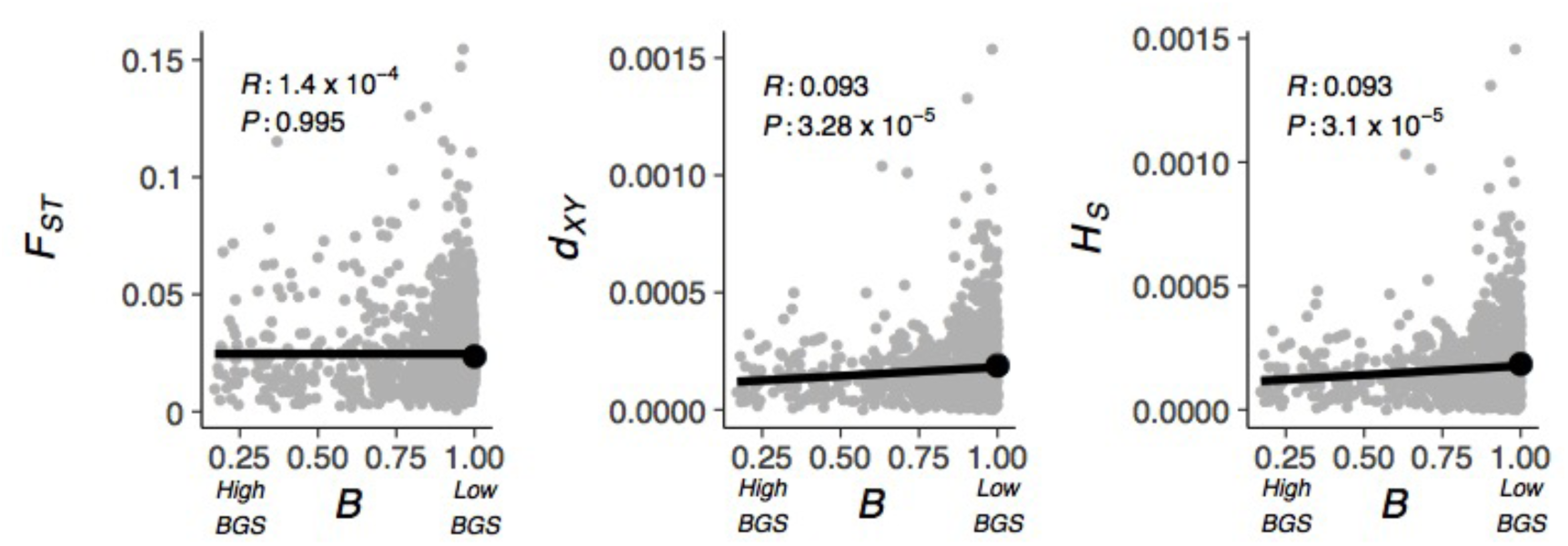
Correlation between *B* and *F*_*ST*_, *H*_*S*_, and *d*_*XY*_ for the last generation (5 × 2*N* generations after the split) of the *Default* treatment. Each grey dot is a single simulation where there is BGS. The large black dot is the mean of the simulations where BGS was artificially turned off. The *P*-values are computed from a Pearson’s correlation test. *P*-values and *R* are computed on the simulations with BGS (grey dots) only.

The *No Migration* treatment is an exception to the other treatments. *F*_*ST*_ is not significantly correlated with *B* at early generations but become slightly correlated as divergence rises to 0.6 and higher. *d*_*XY*_ shows an opposite pattern. *d*_*XY*_ is very significantly correlated with *B* at early generations and seemingly independent of B at the last generation. Note that for both *d*_*XY*_ and *F*_*ST*_, all correlation coefficients are always very small. The largest *R*^2^ observed is *R*^2^=0.0121 in realistic simulations (found for *F*_*ST*_ *No Migration* and for *d*_*XY*_ Large N; Tables S2, S4) and R^2^=0.0256 for the *No Recombination* treatment (found for *d*_*XY*_; Table S4).

As expected, in the *CNC97* simulations, there is a strong difference between simulations with BGS and simulations without BGS for all three statistics (*F*_*ST*_, *d*_*XY*_, and *H*_*S*_) at all generations (Welch’s *t*-tests; all *P* < 2.2×10^−16^; Figure S3).

F_*ST*_ calculated as advised by Weir and Cockerham (1984) was generally less sensitive to BGS than *F*_*ST*_ calculated as an average of ratios (compare tables S2 and S3). Figure S4 illustrates the sensitivity of *F*_*ST* (*average of ratios*)_ in the worst case, the *No Recombination* treatment. This sensitivity is driven largely by rare alleles and goes away when minor alleles below a frequency of 0.05 are excluded.

### F_ST_ outlier tests

The observed false positive rate is relatively close to the α values except for *No Migration* (with and without BGS) and *CNC97* (with BGS). With the exception of treatments *No Migration* and *CNC97*, there is no significant difference in false positive rates between simulations with BGS and those without BGS for α of 0.05 (Figure 3). In the treatment, *No Migration*, the false positive rates between simulations with and without BGS are significantly different for the latter generations. In this treatment, the false positive rate for both simulations with and without BGS are much higher than the α value of 0.05. Results remain very congruent for other α values (results not shown).

**Figure 3:**
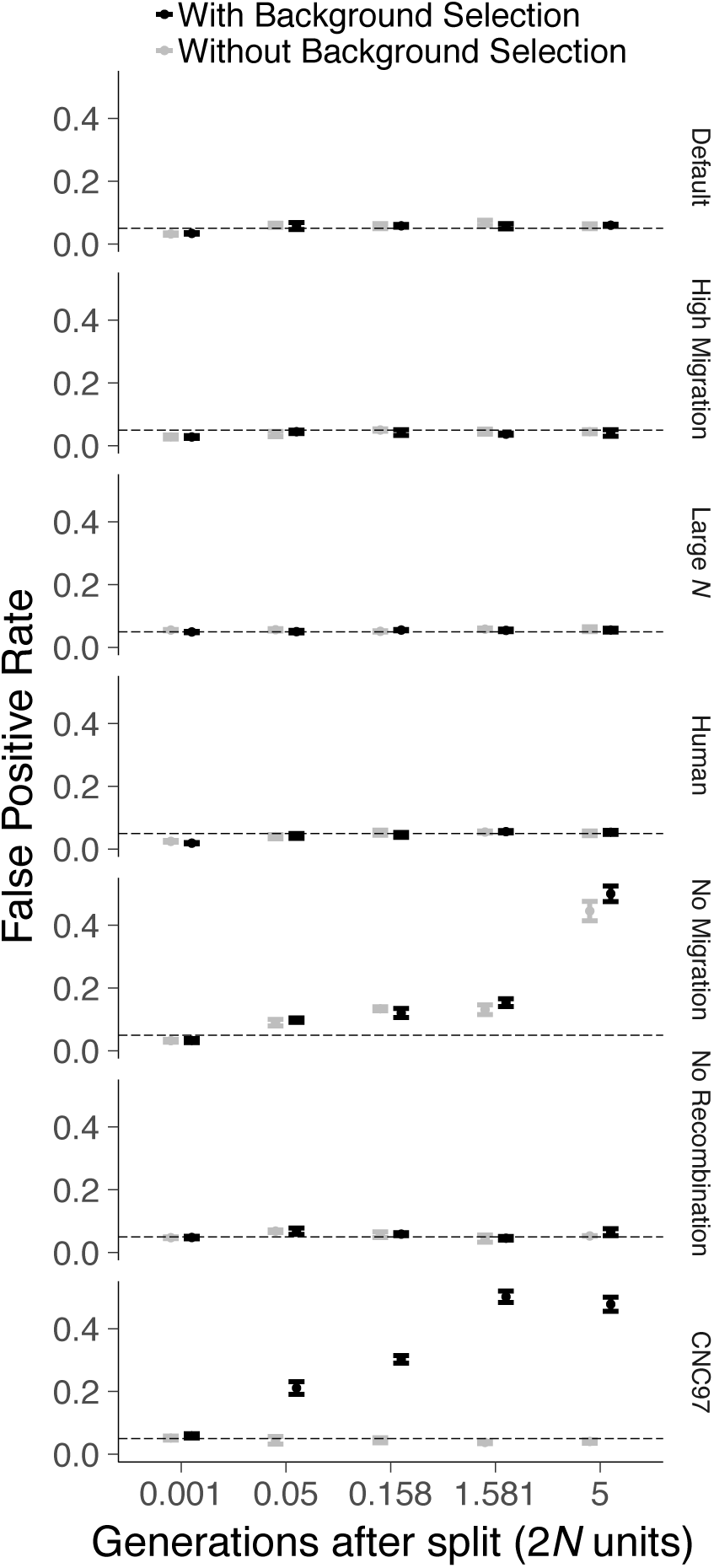
Comparison of false positive rate (FPR) returned by FDist2 between simulations with BGS (black) and without BGS (grey) for all treatments by generation. The significance level is 0.05 and is represented by the horizontal dashed line. Significance based on a Welch’s *t*-test is indicated with stars (2×10^−16^’***’ 0.001 ‘**’ 0.01 ‘*’ 0.05 ‘.’ 0.1 ‘ ’ 1). Significance levels are the same with Wilcoxon tests.

## Discussion

Background selection reduces genetic diversity, both within and among populations, but the effect on *F*_*ST*_ is rather small. In simulations of interconnected populations with realistic parameters, *F*_*ST*_ is insensitive to BGS while the absolute measure of divergence *d*_*XY*_ is affected by BGS. On the other hand, in highly diverged populations unconnected by migration, BGS can have a greater impact on *F*_*ST*_ and a lower impact on *d*_*XY*_. The effects of BGS, when observed, are always very small with *R*^2^ never over 1.1%.

BGS impacts both total and within population genetic diversity. Excluding the treatments *No recombination* and *CNC97*, we observe that simulations with BGS have a genetic diversity (whether *H*_*T*_ or *H*_*S*_; *H*_*T*_ data not shown) 6% to 16% lower than simulations without BGS. Messer & Petrov (2013) simulated a panmictic population, looking at a sequence of similar length inspired from a gene-rich region of the human genome, and reported a similar decrease in genetic diversity. Under the *No Recombination* treatment, this average reduction of genetic diversity due to BGS is 53%. Although empirical estimates are very complex and can hardly disentangle BGS from selective sweeps, our results are also comparable with empirical estimates. Reduction in genetic diversity in humans between regions under high BGS are estimated at 6% according to Cai et al. (2009) or 19-26% according to McVicker et al. (2009). In *Drosophila melanogaster*, where gene density is higher, the reduction in genetic diversity due to BGS is estimated at 36% when using Kim & Stephan (2000)’s methodology and is estimated at 71% reduction using a composite likelihood approach (Elyashiv et al., 2016) and is hence closer to our *No Recombination* treatment than to the other treatments. It is worth noting that, because we were interested in simulating variance among sites in effects of BGS, we only simulated local effects and therefore underestimate the expected genome-wide effect of BGS on *H*_*T*_.

In contrast to measures of heterozygosity, *F*_*ST*_ was generally not significantly correlated with *B*. The only exception is for the *No Migration* treatment, where, after many generations, as the average *F*_*ST*_ becomes very high (*F*_*ST*_ *> 0.5*), we observe a slight, yet significant, negative correlation between the expected effects of BGS, *B*, and *F*_*ST*_ (intense BGS lead to high values of *F*_*ST*_). This highlights that *F*_*ST*_ is not completely insensitive to BGS, but *F*_*ST*_ is largely robust to BGS.

Future research is needed to attempt a theoretical estimate of the genome-wide effect of BGS. Our work has been restricted to the stickleback and human genomes. While these two genomes are good representatives of many cases of eukaryotic genomes, they are not good representatives of more compact genomes such as bacterial genomes or yeasts. Our simulations used randomly mating diploid populations. Non-random mating, selfing, and asexual reproduction could also affect our general conclusion, and potentially strongly increase the effects of BGS on *F*_*ST*_ (Charlesworth *et al.* 1997). We have explored two population sizes, but we could not explore population sizes of the order of a million individuals (like *Drosophila melanogaster*) and still realistically simulate such long stretch of DNA. It is not impossible that a much greater population size or a more complex demography could yield to BGS having a greater effect on *F*_*ST*_ than what we observed here (Torres et al. 2017).

Some have argued that, because BGS reduces the within population diversity, it should lead to high *F*_*ST*_ (Cutter & Payseur, 2013; Cruickshank & Hahn, 2014; Hoban et al., 2016). All else being equal, this statement is correct. However, BGS reduces *H*_*T*_ almost as much as *H*_*S*_ (Figure 4). It is therefore insufficient to consider only one component, and we must consider the ratio of these two quantities captured by the definition of *F*_*ST*_, 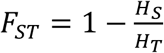. This ratio, as we have shown, appears to be relatively robust to BGS. While genome-wide BGS might eventually be strong enough to cause departures with *F*_*ST*_values, it appears that locus-to-locus variation in the intensity of BGS is not strong enough to have much impact on *F*_*ST*_ as long as populations are not too highly diverged.

**Figure 4:**
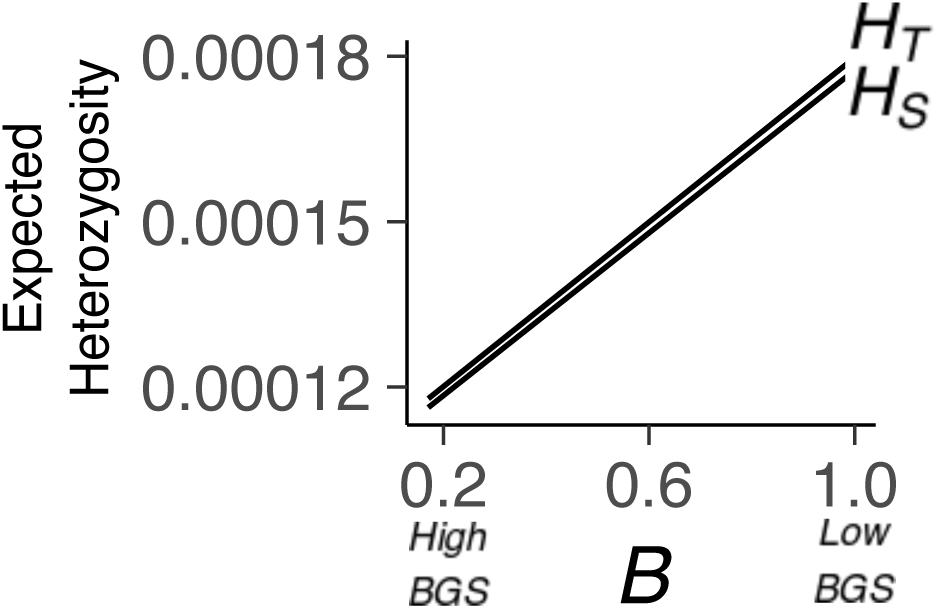
Regressions of total (*H*_*T*_; upper line) and within (*H*_*S*_; lower line) population expected heterozygosity on the coefficient of BGS (*B*) for the last generation of the *Default* treatment. The two regression lines are not exactly parallel with *H*_*S*_ tending to *H*_*T*_ as *B* goes to low values (more intense BGS).

We also investigated the consequences of BGS on the widely-used but imperfect estimator, *F*_*ST* (*average of ratios*)_, for which *F*_*ST*_ measures for each locus are averaged to create a genomic average. It is well known that *F*_*ST* (*average of ratios*)_ is a biased way to average *F*_!”_over several loci (Weir & Cockerham, 1984); however, its usage is relatively common today. In our simulations, *F*_*ST* (*average of ratios*)_ is more affected by BGS than *F*_*ST*_. Interestingly, *F*_*ST* (*average of ratios*)_ is most often higher with weaker BGS. The directionality of this correlation may seem unintuitive at first. To understand this discrepancy, remember that BGS affects the site frequency spectrum; we observed that BGS leads to an excess of loci with low *H*_*T*_ (results not shown but see Charlesworth et al., 1995; see also contrary expectation in Stephan, 2010). Loci associated with very low *H*_*T*_ also have low *F*_*ST*_ (figure S5), a well-known result described by Beaumont and Nichols (1996). As BGS creates an excess of loci with low *H*_*T*_ and loci with low *H*_*T*_ tend to have low *F*_*ST*_, BGS can actually reduce *F*_*ST* (*average of* ratios)_. After filtering out SNPs with a minor allele frequency lower than 5%, most of the correlation between *F*_*ST* (*average of ratios*)_ and *B* is eliminated (Figure S4).

The absolute measure of divergence *d*_*XY*_ is sensitive to BGS. Regions of stronger BGS are associated with low *d*_*XY*_. The effect, although significant, is of relatively small size. The expected *d*_*XY*_ for neutral loci is *d*_*XY*_ *= 4Nµ + 2tµ* (Nei, 1987), where *t* is the time in generation since the populations started to diverge. *4Nµ* is the expected heterozygosity in the ancestral population (before splitting) and *2tµ* is the expected number of mutations fixed over time in either population since the population split. BGS does not affect the rate of fixation of mutations arising after the populations diverged, but BGS affects the expected heterozygosity. Therefore, BGS should affect *d*_*XY*_ by its effect on the expected heterozygosity, and this effect should be greater early in divergence when the 4*N*μ term is large relative to the fixation term. This is consistent with the results of our simulations. This result is in agreement with Vijay et al. (2017) who reported a strong correlation between *H*_*S*_ and *d*_*XY*_ when *F*_*ST*_ is low (*F*_*ST*_ ≈ 0.02), but this correlation breaks down when studying more distantly related populations (*F*_*ST*_ ≈ 0.3).

Interestingly, *d*_*XY*_ becomes less sensitive to BGS when *F*_*ST*_ becomes more sensitive. While our methodology does not allow us to test the efficiency of *d*_*XY*_ in outlier tests, it is possible that *d*_*XY*_ could be used for highly divergent lineages, but not in cases when divergence is relatively low. Cruickshank and Hahn (2014) suggested relying more on *d*_*XY*_ than *F*_*ST*_ for finding highly divergent loci. Their conclusion was based on analysis of a dataset involving highly divergent populations only (*F*_*ST*_ values range from about 0.38 to about 0.8). Based on our simulations, their conclusion is not valid when populations are not highly diverged.

As BGS also leads to a reduction of the number of polymorphic sites, BGS has an even stronger effect on *d*_*XY-SNP*_ than on *d*_*XY*_ (Figure S4). (The measure that we call *d*_*XY*-_ *SNP* is *d*_*xy*_ improperly calculated based only on polymorphic sites, as is done in some software packages.) This result highlights the importance of not blindly trusting the output of a given software package.

Outside the effect of BGS on *N*_*e*_, there are at least two other possible factors that can potentially affect the correlation between *B* and *µ*: the effect of deleterious mutations on the effective migration rate and the auto-correlation of µ. Because most deleterious mutations are recessive (García-Dorado and Caballero, 2000; Peters et al., 2003; Shaw & Chang, 2006), the offspring of migrants, who enjoy an increased heterozygosity compared to local individuals, will be at a selective advantage. The presence of deleterious mutations therefore lead to an increase in the effective migration rate (Ingvarsson & Whitlock, 2000). This increases the effective migration rate and hence, leads to a decrease in *F*_*ST*_.

As mutation rate is auto-correlated throughout the genome, neutral sequences closely linked to sequences that frequently receive deleterious mutation are also likely to experience frequent neutral mutations. As a high mutation rate leads to low *F* values (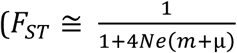, Wright 1943), autocorrelation in mutation rate may also impact the correlation between *B* and *F*_*ST*_. This effect is likely to be negligible as long as *m* >> µ.

Recently, evidence of a correlation between recombination rate and *F*_*ST*_ has been interpreted as likely being caused by deleterious mutations rather than positive selection, whether the divergence between populations is very high (e.g. Cruickshank & Hahn, 2014), moderately high (Vijay et al., 2017) or moderately low (Torres et al., 2017). Here we showed the BGS is unlikely to explain all of these correlations. It can be hypothesized that positive selection (selective sweeps and local adaptation) could be the main cause of this correlation.

McVicker et al. (2009) attempted an estimation of *B* values in the human genome (see also Elyashiv et al., 2016). They did so using equations from Nordborg et al. (1996). As there is little knowledge about the strength of selection throughout the genome, to our understanding, this estimation of *B* values should be highly influenced by the effects of beneficial mutations as well as deleterious mutations. Torres et al. (2017) reused this dataset and found a slight association between *B* and *F*_*ST*_ among human lineages. It is plausible that this correlation between *B* and *F*_*ST*_ could be driven by positive selection rather than by deleterious mutations.

Our FDist2 analysis shows that the false positive rate does not differ in simulations with BGS or without BGS. The only exceptions concern the unrealistic *CNC97* treatment and the *No Migration* treatment after many generations (Figure 3). The average *F*_*ST*_ at the last generation of the *No Migration* treatment is greater than 0.8. With such high *F*_*ST*_ both the simulation without BGS and with BGS lead to very high false positive rates (0.45 without BGS and 0.5 with BGS). This difference in false positive rates is significant, but the observed high false positive rate should make it clear that for such highly diverged populations, *F*_*ST*_ outlier tests are not recommended in general.

Many authors (e.g. Nachman & Payseur, 2012; Cutter & Payseur, 2013; McGee et al., 2015; Yeaman, 2015; Whitlock & Lotterhos, 2015; Brousseau et al., 2016; Picq et al., 2016; Payseur & Rieseberg, 2016; Hoban et al., 2016) have raised concerns that BGS can strongly reduce our ability to detect the genomic signature of local adaptation. Our analysis shows that BGS is not a strong confounding factor to *F*_*ST*_ outlier tests of populations that are not too highly diverged.

## Acknowledgments

Many thanks to Loren H. Rieseberg, Sarah P. Otto and Amy L. Angert for their help in discussing the design of the project and for feedback. Thanks to Sarah P. Otto, Darren E. Irwin, and Bret A. Payseur for helpful comments on the manuscript. We also thank Yaniv Brandvain and Tom Booker for their feedback and Marius Roesti for help with the Ensembl-retrieved gene annotations.

---

**Table S1:**
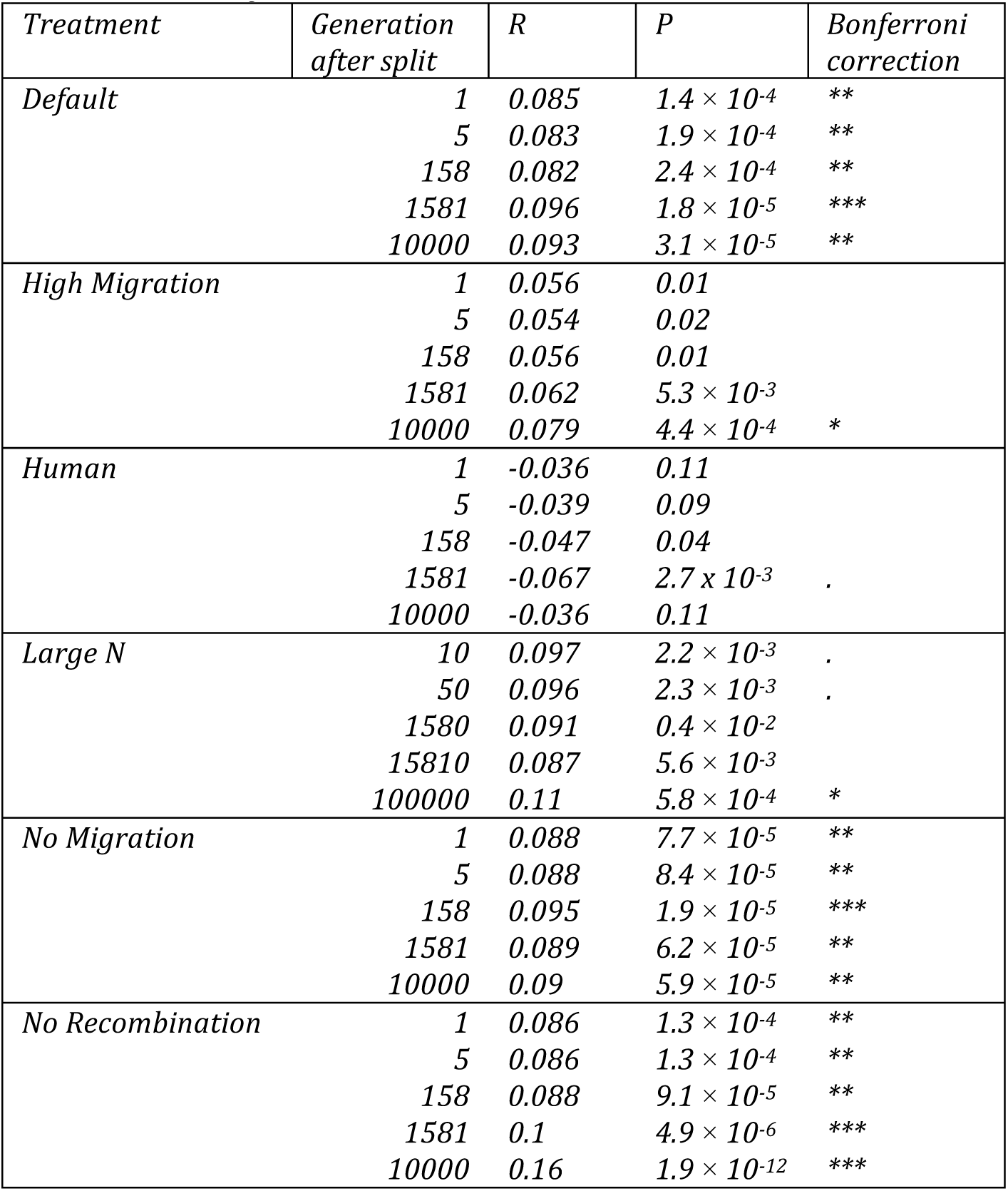
Pearson’s correlation tests for the association between the coefficient of background selection (*B*) and *H*_*S*_. *P*-values and *R* are computed on the simulations with BGS only.

| *Treatment* | *Generation*<br>*after split* | *R* | *P* | *Bonferroni*<br>*correction* |
| --- | --- | --- | --- | --- |
| *Default* | 1 | 0.085 | $1.4 \times 10^{-4}$ | ** |
| | 5 | 0.083 | $1.9 \times 10^{-4}$ | ** |
| | 158 | 0.082 | $2.4 \times 10^{-4}$ | ** |
| | 1581 | 0.096 | $1.8 \times 10^{-5}$ | *** |
| | 10000 | 0.093 | $3.1 \times 10^{-5}$ | ** |
| *High Migration* | 1 | 0.056 | 0.01 |  |
|  | 5 | 0.054 | 0.02 |  |
|  | 158 | 0.056 | 0.01 |  |
| | 1581 | 0.062 | $5.3 \times 10^{-3}$ | |
| | 10000 | 0.079 | $4.4 \times 10^{-4}$ | * |
| *Human* | 1 | -0.036 | 0.11 |  |
|  | 5 | -0.039 | 0.09 |  |
|  | 158 | -0.047 | 0.04 |  |
| | 1581 | -0.067 | $2.7 \times 10^{-3}$ | . |
|  | 10000 | -0.036 | 0.11 |  |
| *Large N* | 10 | 0.097 | $2.2 \times 10^{-3}$ | . |
| | 50 | 0.096 | $2.3 \times 10^{-3}$ | . |
| | 1580 | 0.091 | $0.4 \times 10^{-2}$ | |
| | 15810 | 0.087 | $5.6 \times 10^{-3}$ | |
| | 100000 | 0.11 | $5.8 \times 10^{-4}$ | * |
| *No Migration* | 1 | 0.088 | $7.7 \times 10^{-5}$ | ** |
| | 5 | 0.088 | $8.4 \times 10^{-5}$ | ** |
| | 158 | 0.095 | $1.9 \times 10^{-5}$ | *** |
| | 1581 | 0.089 | $6.2 \times 10^{-5}$ | ** |
| | 10000 | 0.09 | $5.9 \times 10^{-5}$ | ** |
| *No Recombination* | 1 | 0.086 | $1.3 \times 10^{-4}$ | ** |
| | 5 | 0.086 | $1.3 \times 10^{-4}$ | ** |
| | 158 | 0.088 | $9.1 \times 10^{-5}$ | ** |
| | 1581 | 0.1 | $4.9 \times 10^{-6}$ | *** |
| | 10000 | 0.16 | $1.9 \times 10^{-12}$ | *** |

**Table S2:**
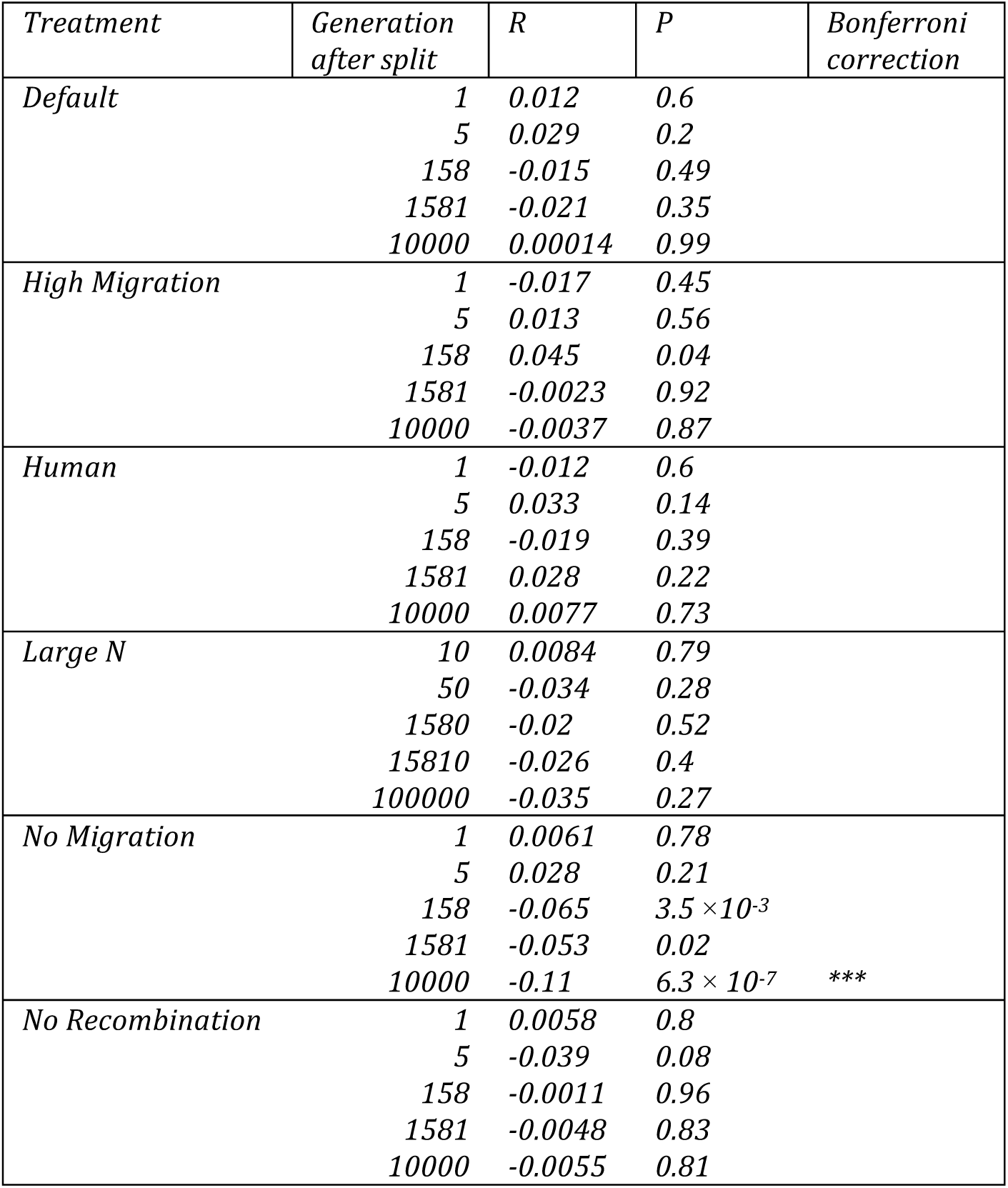
Pearson’s correlation tests for the association between the coefficient of background selection (*B*) and *F*_*ST*_. *P*-values and *R* are computed on the simulations with BGS only.

**Table S3:**
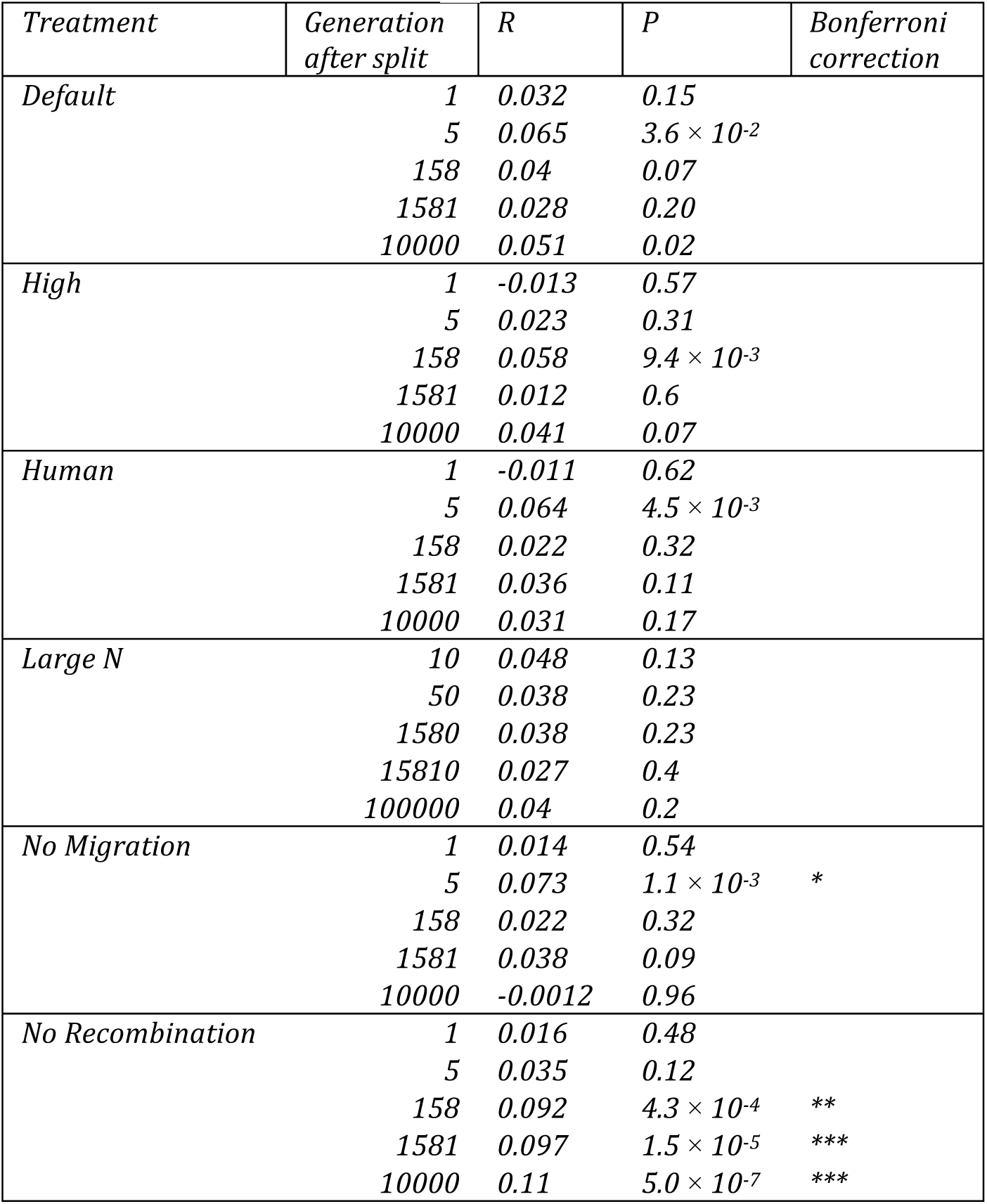
Pearson’s correlation tests for the association between the coefficient of background selection (*B*) and *F*_*ST* (average of ratios)_. *P*-values and *R* are computed on the simulations with BGS only.

| *Treatment* | *Generation*<br>*after split* | *R* | *P* | *Bonferroni*<br>*correction* |
| --- | --- | --- | --- | --- |
| *Default* | 1 | 0.032 | 0.15 |  |
| | 5 | 0.065 | $3.6 \times 10^{-2}$ | |
|  | 158 | 0.04 | 0.07 |  |
|  | 1581 | 0.028 | 0.20 |  |
|  | 10000 | 0.051 | 0.02 |  |
| *High* | 1 | -0.013 | 0.57 |  |
|  | 5 | 0.023 | 0.31 |  |
| | 158 | 0.058 | $9.4 \times 10^{-3}$ | |
|  | 1581 | 0.012 | 0.6 |  |
|  | 10000 | 0.041 | 0.07 |  |
| *Human* | 1 | -0.011 | 0.62 |  |
| | 5 | 0.064 | $4.5 \times 10^{-3}$ | |
|  | 158 | 0.022 | 0.32 |  |
|  | 1581 | 0.036 | 0.11 |  |
|  | 10000 | 0.031 | 0.17 |  |
| *Large N* | 10 | 0.048 | 0.13 |  |
|  | 50 | 0.038 | 0.23 |  |
|  | 1580 | 0.038 | 0.23 |  |
|  | 15810 | 0.027 | 0.4 |  |
|  | 100000 | 0.04 | 0.2 |  |
| *No Migration* | 1 | 0.014 | 0.54 |  |
| | 5 | 0.073 | $1.1 \times 10^{-3}$ | * |
|  | 158 | 0.022 | 0.32 |  |
|  | 1581 | 0.038 | 0.09 |  |
|  | 10000 | -0.0012 | 0.96 |  |
| *No Recombination* | 1 | 0.016 | 0.48 |  |
|  | 5 | 0.035 | 0.12 |  |
| | 158 | 0.092 | $4.3 \times 10^{-4}$ | ** |
| | 1581 | 0.097 | $1.5 \times 10^{-5}$ | *** |
| | 10000 | 0.11 | $5.0 \times 10^{-7}$ | *** |

**Table S4:** Pearson’s correlation tests for the association between the coefficient of background selection (*B*) and *d*_*XY*_. *P*-values and *R* are computed on the simulations with BGS only.

| *Treatment* | *Generation*<br>*after split* | *R* | *P* | *Bonferroni*<br>*correction* |
| --- | --- | --- | --- | --- |
| *Default* | 1 | 0.085 | $1.35 \times 10^{-4}$ | ** |
| | 5 | 0.084 | $1.35 \times 10^{-4}$ | ** |
| | 158 | 0.082 | $2.46 \times 10^{-4}$ | ** |
| | 1581 | 0.095 | $2.12 \times 10^{-5}$ | *** |
| | 10000 | 0.093 | $3.28 \times 10^{-5}$ | ** |
| *High Migration* | 1 | 0.056 | 0.01 |  |
|  | 5 | 0.054 | 0.02 |  |
|  | 158 | 0.056 | 0.01 |  |
| | 1581 | 0.062 | $5.26 \times 10^{-3}$ | |
| | 10000 | 0.079 | $4.36 \times 10^{-4}$ | * |
| *Human* | 1 | -0.036 | 0.11 |  |
|  | 5 | -0.037 | 0.10 |  |
|  | 158 | -0.047 | 0.03 |  |
| | 1581 | -0.067 | $3.0 \times 10^{-3}$ | |
|  | 10000 | -0.036 | 0.11 |  |
| *Large N* | 10 | 0.097 | $2.2 \times 10^{-3}$ | . |
| | 50 | 0.096 | $2.3 \times 10^{-3}$ | . |
| | 1580 | 0.091 | $4.0 \times 10^{-3}$ | |
| | 15810 | 0.087 | $5.7 \times 10^{-3}$ | |
| | 100000 | 0.11 | $5.8 \times 10^{-4}$ | * |
| *No Migration* | 1 | 0.088 | $7.6 \times 10^{-5}$ | ** |
| | 5 | 0.089 | $6.7 \times 10^{-5}$ | ** |
| | 158 | 0.086 | $1.2 \times 10^{-4}$ | ** |
|  | 1581 | 0.052 | 0.02 |  |
|  | 10000 | 0.029 | 0.20 |  |
| *No Recombination* | 1 | 0.086 | $1.3 \times 10^{-4}$ | ** |
| | 5 | 0.085 | $1.6 \times 10^{-4}$ | ** |
| | 158 | 0.088 | $8.8 \times 10^{-5}$ | ** |
| | 1581 | 0.1 | $4.9 \times 10^{-5}$ | *** |
| | 10000 | 0.16 | $2.3 \times 10^{-12}$ | *** |

**Table S5:**
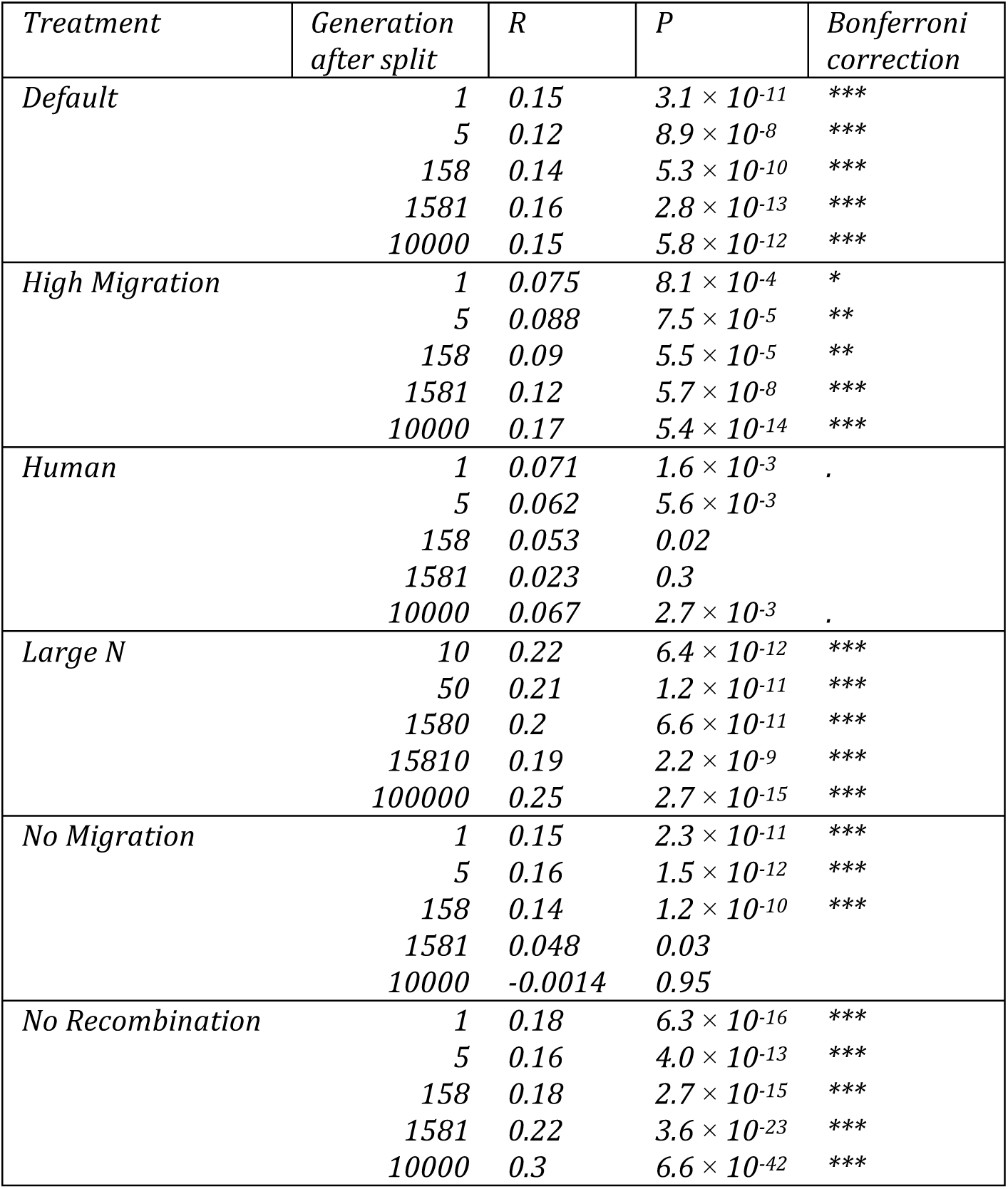
Pearson’s correlation tests for the association between the coefficient of background selection (*B*) and *d*_*XY*-SNP_. *P*-values and *R* are computed on the simulations with BGS only.

**Figure S1:**
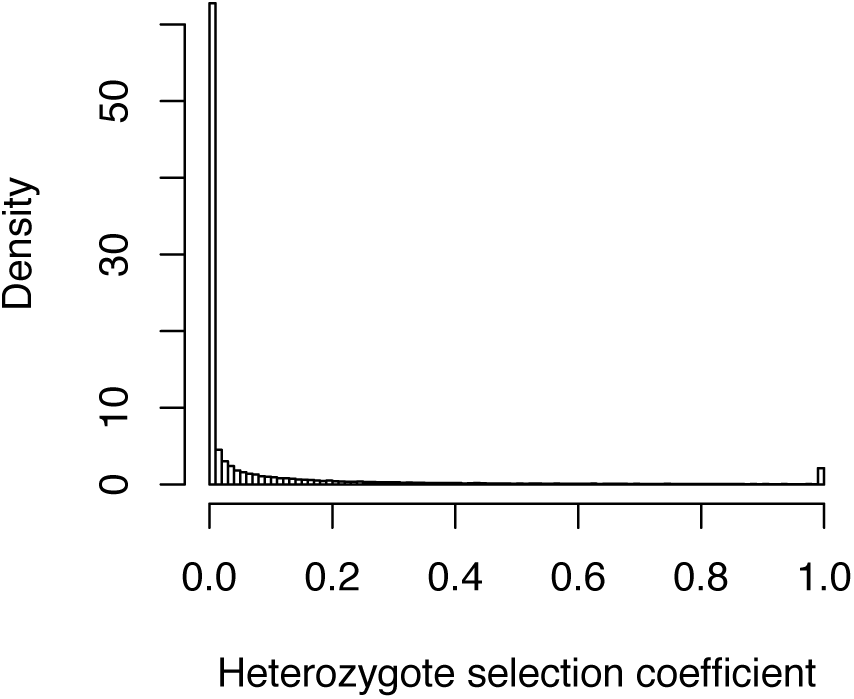
Overall distribution of selection coefficient in the heterozygotes. There are 2% lethal mutations, and the average selection coefficient of the non-lethal mutations is approximately 0.07.

**Figure S2:**
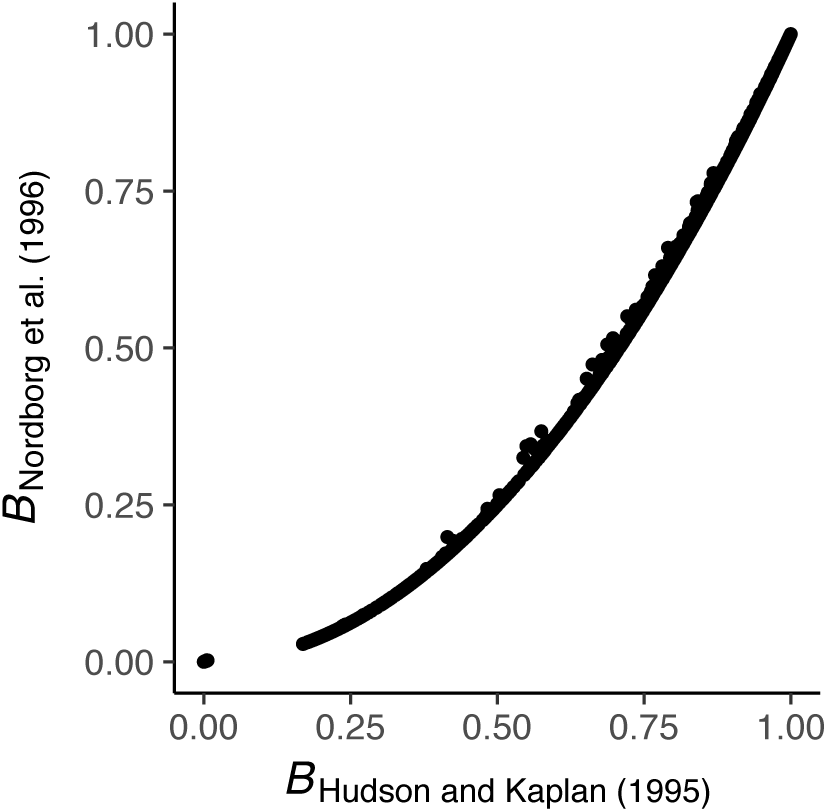
Relationship between *B* values computed for all simulations using the stickleback genome using the methods of Hudson and Kaplan (1996) and Nordborg et al. (1997).

**Figure S3:**
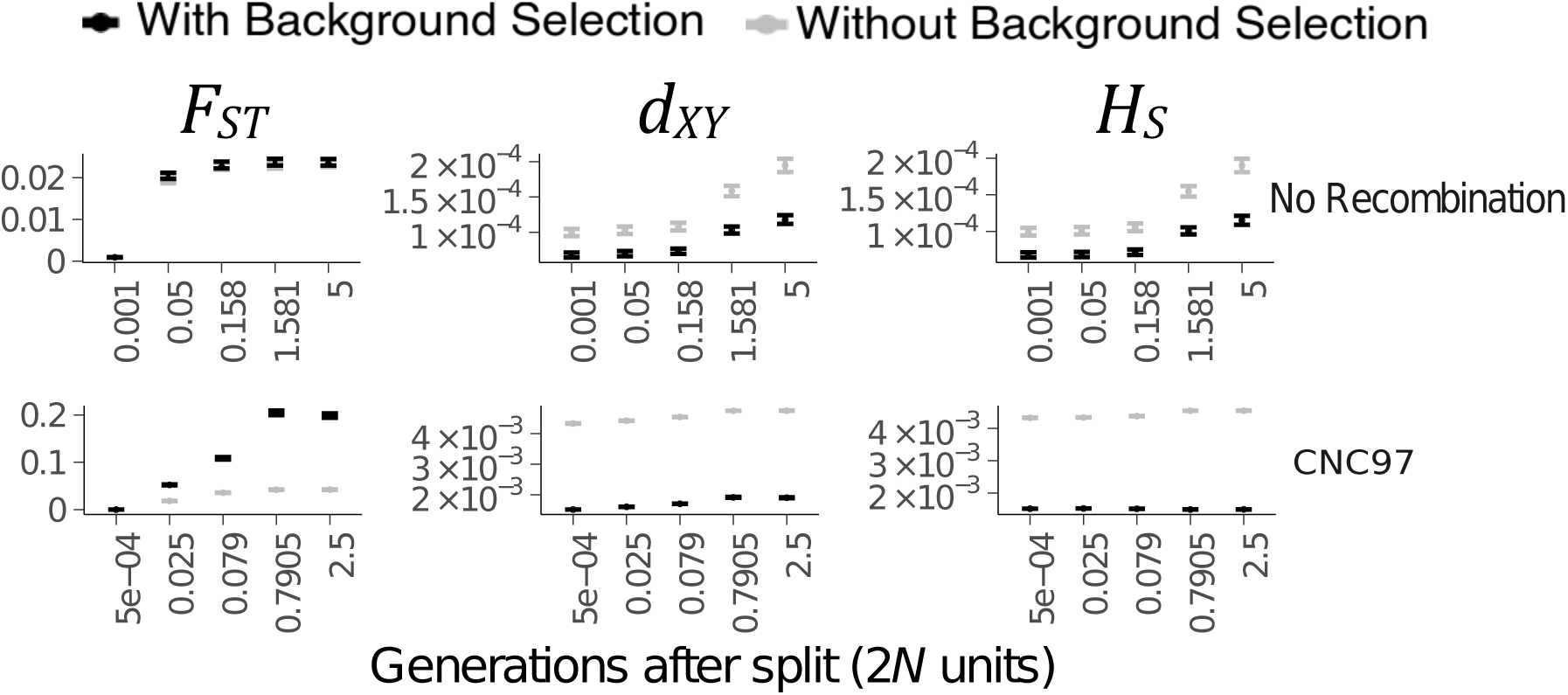
Comparisons of means *F*_*ST*_ (left column), *d*_*XY*_ (central column), and *H*_*S*_ (right column) between simulations with (black) and without (grey) BGS for all unrealistic treatments. Realistic treatments (*Default, No Migration, High Migration, Human* and *Large N*) are in Figure 1 in the main text. Error bars are 95% confidence intervals.

**Figure S4:**
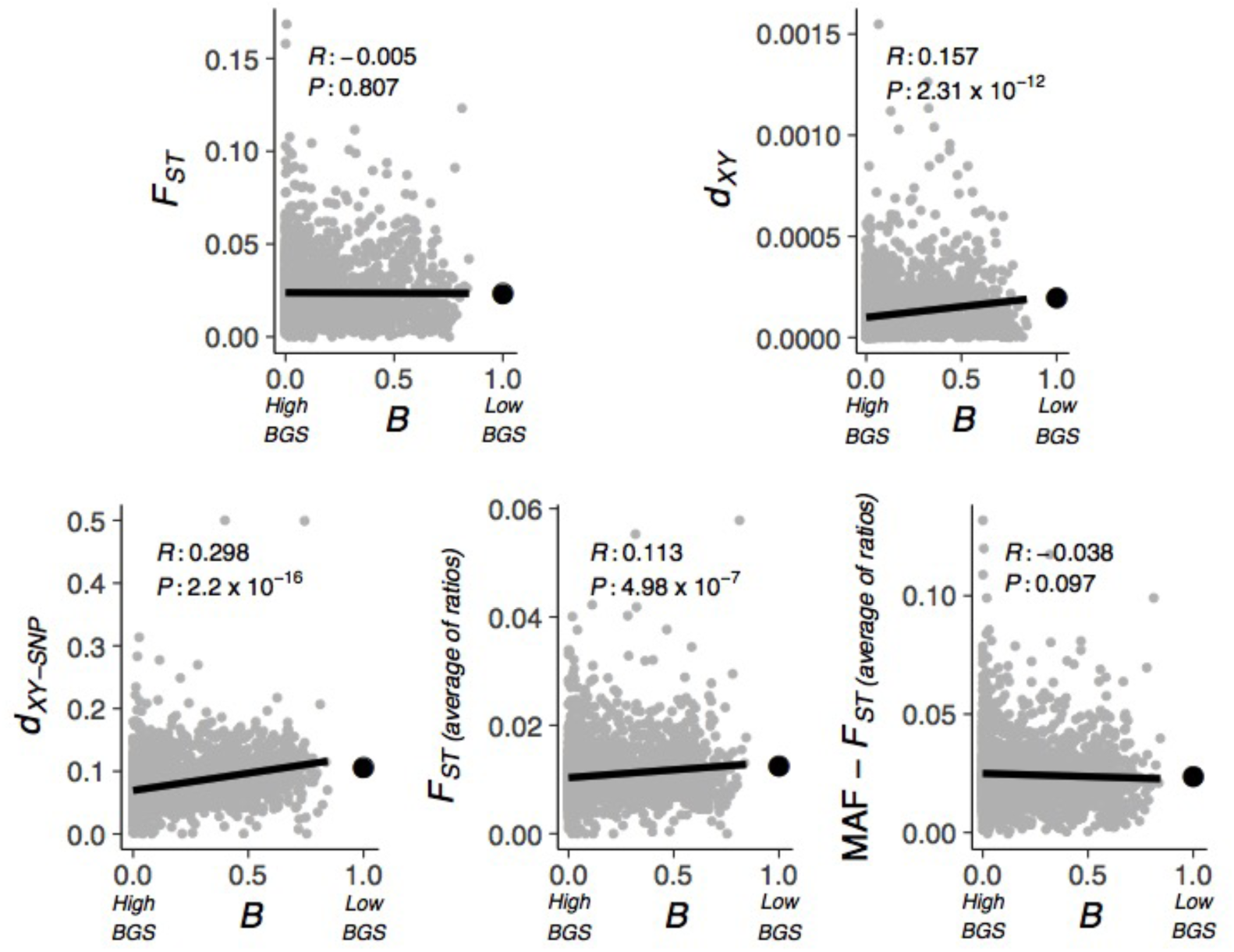
Correlations between *B* and *F*_*ST*_, *d*_*xy*_, *F*_*ST*_ (average of ratios), *d*_*xy*_-SNP, and *F*_*ST*_ (average of ratios) after removing all loci that have minor allele frequency (MAF) lower than 0.05 (called MAF - *F*_*ST*_ (average of ratios)) for the treatment *No Recombination* only at the last generation (5 x 2*N* generations after the split). Each grey dot is a single simulation with BGS. The large black dot is the mean of all simulations without BGS. The *P*-values are computed from a Pearson’s correlation test. *P*-values and *R* are computed on the simulations with BGS (grey dots) only.

**Figure S5:**
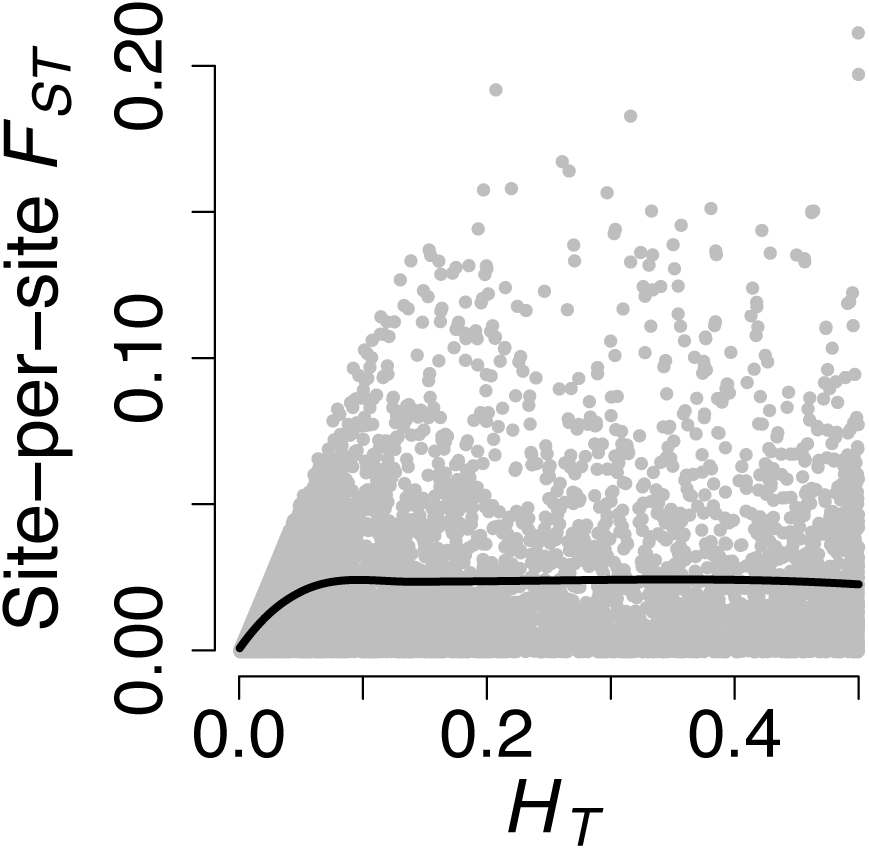
Relationship between total heterozygosity *H*_*T*_ and *F*_*ST*_ on a site per site basis. Each dot represents a single site. The black line is a Local Polynomial Regression (LOESS). Data is a random subset from the *Default* treatment at the last generation.

